# Genomic comparison of the temperate coral *Astrangia poculata* with tropical corals yields insights into winter quiescence, innate immunity, and sexual reproduction

**DOI:** 10.1101/2023.09.22.558704

**Authors:** Kathryn H. Stankiewicz, Nadège Guiglielmoni, Sheila A. Kitchen, Jean-François Flot, Katie L. Barott, Sarah W. Davies, John R. Finnerty, Sean P. Grace, Leslie S. Kaufman, Hollie M. Putnam, Randi D. Rotjan, Koty H. Sharp, Iliana B. Baums

## Abstract

Facultatively symbiotic corals provide important experimental models to explore the establishment, maintenance, and breakdown of the mutualism between corals and members of the algal family Symbiodiniaceae. The temperate coral *Astrangia poculata* is one such model as it is not only facultatively symbiotic, but also occurs across a broad temperature and latitudinal gradient. Here, we report the *de novo* chromosome-scale assembly and annotation of the *A. poculata* genome. Though widespread segmental/tandem duplications of genomic regions were detected, we did not find strong evidence of a whole genome duplication (WGD) event. Comparison of the gene arrangement between *A. poculata* and the tropical coral *Acropora millepora* revealed 56.38% of the orthologous genes were conserved in syntenic blocks despite ∼415 million years of divergence. Gene families related to sperm hyperactivation and innate immunity, including lectins, were found to contain more genes in *A. millepora* relative to *A. poculata*. Sperm hyperactivation in *A. millepora* is expected given the extreme requirements of gamete competition during mass spawning events in tropical corals, while lectins are important in the establishment of coral-algal symbiosis. By contrast, gene families involved in sleep promotion, feeding suppression, and circadian sleep/wake cycle processes were expanded in *A. poculata*. These expanded gene families may play a role in *A. poculata*’s ability to enter a dormancy-like state (“winter quiescence”) to survive freezing temperatures at the northern edges of the species’ range.

## Introduction

Anthozoa, the largest class within the phylum Cnidaria, includes some of the most ecologically important and oldest clades of marine metazoans, estimated to have evolved as early as 771 million years ago (Mya) (McFadden et al., 2021). Among these are corals, a diverse group of colonial organisms that can form a symbiotic association with algae of the family Symbiodiniaceae in the shallow water of the tropic and temperate zones (LaJeunesse et al., 2018). Stony corals of the order Scleractinia contain the engineers of reef ecosystems and are generally divided into two major clades, Complexa and Robusta, which diverged approximately 415 Mya (Kitahara, Cairns, Stolarski, Blair, & Miller, 2010; Romano & Palumbi, 1996; Stolarski et al., 2011). Over the past several decades, tropical coral species have undergone mass mortality due to their sensitivity to bleaching in the face of anthropogenic climate change (Bellwood, Hughes, Folke, & Nyström, 2004; DeCarlo et al., 2017; Hughes et al., 2017). Understanding differences between species of variable temperature tolerance and adaptive strategies has become a focus for conservation efforts. In the “robust” clade, the temperate coral *Astrangia poculata* (the Northern Star Coral) has recently emerged as a model system for these comparisons due to its facultative symbiosis and temperature tolerance ranging from near freezing to 24°C in Narragansett Bay, the northern part of its distribution (Jacques, Marshall, & Pilson, 1983).

*Astrangia poculata,* like many other shallow water corals, hosts a photosynthesizing algal symbiont, *Breviolum psygmophilum* (Lajeunesse, Parkinson, & Reimer, 2012). The symbiosis is facultative, with a gradient of symbiont density existing among individual polyps within a single colony of *A. poculata* and between sympatric colonies (Dimond & Carrington, 2007) as revealed by an aposymbiotic ‘white’ appearance, nearly or entirely devoid of symbionts, or a symbiotic ‘brown’ appearance (Figure 1A). Unlike many other shallow water coral species, *A. poculata* occurs across a broad temperature and latitudinal gradient from southern Massachusetts to the Gulf of Mexico (Cummings, 1983; Dimond & Carrington, 2007; Peters, 1988; Dimond et al 2013). At the northernmost parts of its range, during winter months high in the intertidal zone, *A. poculata* experiences a quiescence characterized by a dormancy state of reduced feeding and growth (Figure 1B) (Grace, 2017; Jacques et al., 1983). This appears to be an adaptation of *A. poculata* to the intertidal zone, where it is vulnerable to desiccation, predation, and extreme shifts in salinity and temperature. *A. poculata’s* hardiness combined with the ability of researchers to experimentally isolate the contributions of the host and the symbiont in aposymbiotic and symbiotic colonies has made this species particularly interesting for the study of coral symbiosis in the face of climate change.

**Figure 1:**
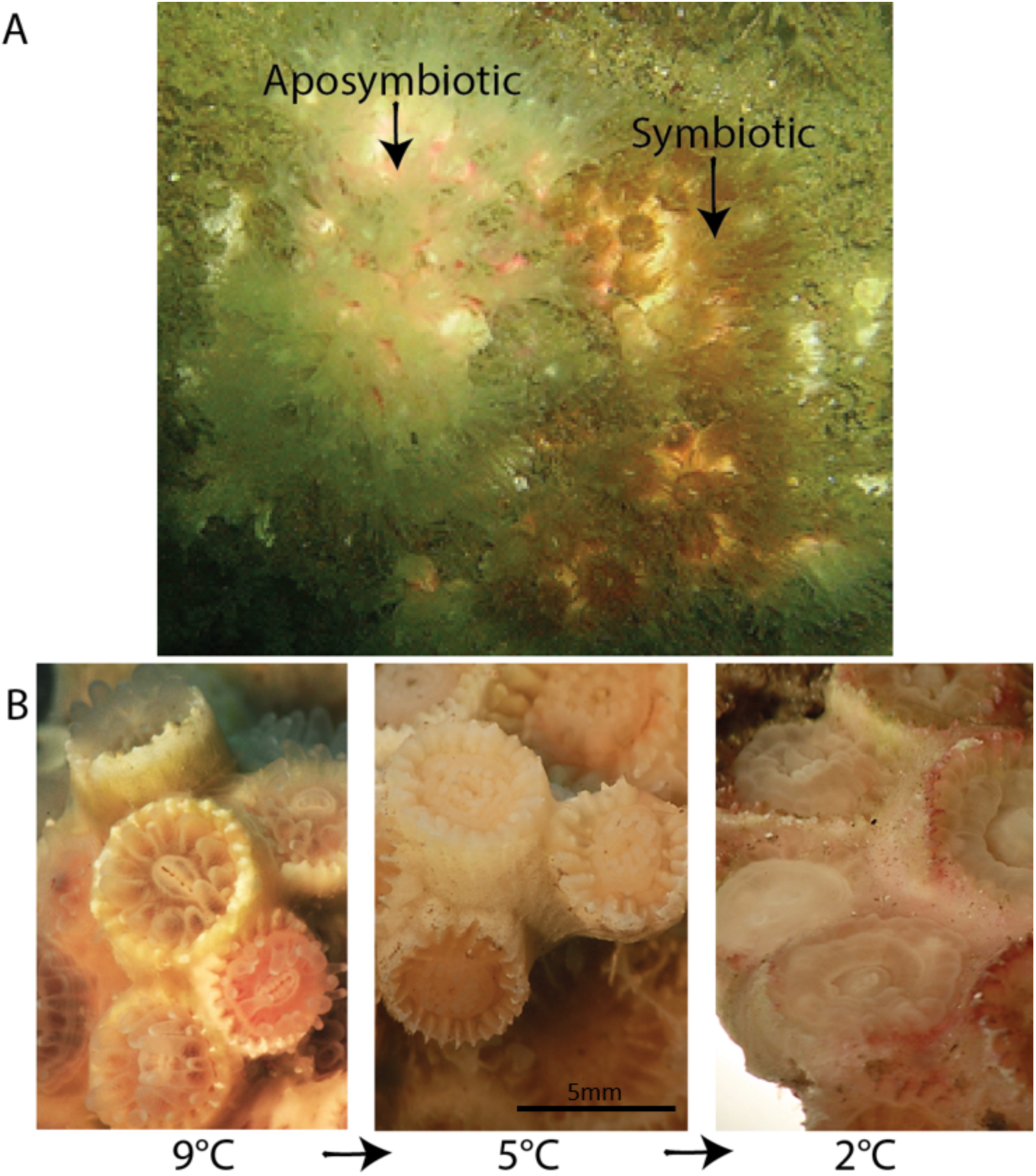
Underwater photograph of *Astrangia poculata* colony with (**A**) extended tentacles exhibiting aposymbiotic (white appearance, left) and symbiotic (brown appearance, right) states and (**B)** transitioning into quiescence from 9°C (pre-dormancy state) to 5°C (transitionary state; retracting tentacles) to 2°C (dormant; tentacles fully retracted, oral disk puffed out).

Previous work found signatures of adaptation in thermal tolerance of *A. poculata* across the species’ range, with a cold-adapted population exhibiting higher metabolic rates than warm adapted populations across several temperature treatments (Aichelman, Zimmerman, & Barshis, 2019). Symbiotic and aposymbiotic *A. poculata* have been shown to respond differently to warm and cold temperatures at both the organismal and transcriptomic levels. Burmester et al. (2017) found that symbiotic colonies significantly outperformed aposymbiotic colonies at 9, 18, and 24°C with respect to wound healing, while Wuitchik et al. (2021) found that aposymbiotic colonies responded more strongly transcriptionally to a cold exposure than to a heat exposure treatment. In a separate thermal stress experiment, the algal endosymbiont *Breviolum psygmophilum* responded more strongly than the *A. poculata* host to chronic heat stress (Chan et al., 2021). Further, season, rather than symbiont state, was shown to drive the structure of the microbiome in *A. poculata* (Sharp, Pratte, Kerwin, Rotjan, & Stewart, 2017). Thus, across multiple scales of measurement, thermal variation, particularly exposure to cold temperatures, appears to play an important role in the biology and ecology of this coral. However, the evolutionary roots and the genomic mechanisms driving the response to environmental change remain unclear for *A. poculata,* and for many other corals.

The use of comparative genomics has shed light on the evolution of basal metazoans. For example, studies have suggested the potential roles of whole-genome duplication (Mao & Satoh, 2019), horizontal gene transfer (Bhattacharya et al., 2016), and *de novo* biosynthesis pathways (Ying et al., 2018) in coral evolutionary trajectories. Many of these comparisons have focused on complex versus robust lineages, for which there exists a deep evolutionary split based on molecular and phylogenetic evidence (Kitahara et al., 2010; Romano & Palumbi, 1996; Stolarski et al., 2011). However, it is not well understood how phylogenetically widespread these genomic traits may be, and the inclusion of other corals representing a wider array of ecological niches and adaptive strategies is warranted, which requires additional genomic resources.

While over the past several years there has been an increase in the availability of genomic resources for cnidarians, currently few of them are at chromosome-scale (however, see Fuller et al., 2020; Hu et al., 2020; Stephens et al., 2022) and none represent a facultatively symbiotic, temperate coral. To fill this gap, we present here the chromosome-scale assembly of *A. poculata*. The assembly is among the most contiguous and complete of available coral genome assemblies available to date. We characterized the structure and content of the *A. poculata* genome to investigate potential genomic drivers underlying this species’ unique temperature tolerance and flexible symbiotic state. Our aims were to 1) produce a high-quality reference genome for *A. poculata*, 2) use this genome assembly to explore several potential genomic mechanisms that may contribute to the unique plasticity of the species, and 3) characterize the degree of similarity of the gene repertoire and genome organization among *A. poculata* and other corals.

## Methods

### Genome sequencing and *de novo* assembly

An aposymbiotic colony was collected from Fort Wetherill State Park in Jamestown, Rhode Island, USA in October of 2017. To avoid sequencing the symbiont, a colony with a minimal density of *B. psygmophilum* in its tissue was selected based on the colony’s white appearance.

Sequencing included two Illumina datasets of paired-end 150 base-pair (bp) reads: one with 414 million reads and an estimated average insert size of 395 bp, and the second with 235 million reads and an estimated average insert size of 484 bp. Three Hi-C libraries were produced with 198 million, 266 million, and 257 million of 150 bp paired-end reads each. Library preparation and sequencing of the Illumina and Hi-C data were carried out by Dovetail Genomics (https://dovetailgenomics.com/). All Illumina and Hi-C reads were trimmed using Cutadapt v2.9 with default settings (Martin, 2011).

In preparation for Oxford Nanopore sequencing, extracted genomic DNA was purified with AMPure XP Beads to improve the purity. Short fragments were discarded using Circulomics Short Reads Eliminator XS (Circulomics, Pacific Biosciences). The library was prepared using the Nanopore Ligation Sequencing Kit LSK109, starting with 2.1 μg of DNA, and yielded 1.4 μg of library. The final library was sequenced with a MinION on a R9.4 flowcell with fast basecalling (Jain, Olsen, Paten, & Akeson, 2016). The flowcell was washed and reloaded three times (281 ng of DNA for the first load, 187 ng for subsequent loads). A total output of 6.79 Gigabases (Gb) was obtained with an N_50_ of 18 kilobases (kb) and an N_90_ of 5 kb. The reads were trimmed of the adaptors with Porechop (available at https://github.com/rrwick/Porechop), using default parameters. After trimming, the dataset reached 6.77 Gb.

Several assembly strategies were compared to select the most robust approach. Assemblers that were investigated include Ra (Vaser & Šikić 2019), Raven (Vaser & Šikić 2019), Flye (Kolmogorov, Yuan, Lin, & Pevzner, 2019), Canu (Koren et al., 2017), and wtdbg2 (Ruan & Li, 2020). Ultimately, the genome was assembled using wtdbg2 v2.5 under default settings. Haplotigs were purged using purge_haplotigs v1.1.1 (Roach, Schmidt, & Borneman, 2018) with default settings (following the recommendations in Guiglielmoni et al. (2021)) with both Illumina datasets mapped using bowtie2 v2.3.5.1 (Langmead & Salzberg, 2012). Polishing was done using HyPo v1.0.3 (Kundu, Casey, & Sung, 2019). Hi-C reads were mapped to the assembly and processed using bowtie2 v2.3.5.1 and hicstuff v2.3.0 (Matthey-Doret et al., 2020) with the parameters --enzyme DpnII --iterative --aligner bowtie2. The draft assembly was then scaffolded using instaGRAAL v0.1.6 no-opengl branch (Baudry et al., 2020), with default parameters (--levels 4, --cycles 100, --coverage-std 1, --neighborhood 5). The output was then refined using the module instaGRAAL-polish. BUSCO v4.1.4 was used to assess the completeness of the assembly (Simão, Waterhouse, Ioannidis, Kriventseva, & Zdobnov, 2015) against the Metazoa odb9 lineage. The genome size was estimated with the Illumina dataset of 235 million reads and the module kmercount.sh from BBmap v38.79 (Bushnell, 2014). The circularized mitochondrial genome was assembled from the Illumina reads using NOVOPlasty v 2.7.2 (Dierckxsens, Mardulyn, & Smits, 2016) with the publicly available *Acropora digitifera* mitochondrial genome as a reference (GenBank: KF448535.1). Our 14 chromosome-scale scaffolds were named Ap1, Ap2… Ap14 on the basis of their homologies with *Acropora millepora* chromosome-scale scaffolds bearing the same number (in the absence of a published karyotype for *Astrangia poculata*).

### Transcriptome assembly

RNA sequencing data from Chan et al. (2021) were used to construct a *de novo* transcriptome. First, to limit the potential for symbiont contamination, only reads for aposymbiotic individuals were used for the host transcriptome assembly. Reads were trimmed using Cutadapt v 3.4 (Martin, 2011) with a quality cut-off of 15 and a minimum read length of 50 nucleotides. The trimmed reads were then assembled into transcripts using rnaSPades v 3.12.0 (Bushmanova, Antipov, Lapidus, & Prjibelski, 2019). To further reduce the possibility of contamination from the algal endosymbiont, a custom database of Symbiodiniaceae protein sequences was assembled that included the following species: *Cladocopium goreaui* (H. Liu et al., 2018), *Breviolum psygmophilum* (Parkinson et al., 2016), *Breviolum minutum* (Shoguchi et al., 2013), *Fugacium kawaguti* (H. Liu et al., 2018), *Symbiodinium fitti* (Reich et al., 2021), *Symbiodinium microadriaticum* (Aranda et al., 2016), and *Symbiodinium tridacnidorum* (González-Pech et al., 2021). A BLAST nucleotide-to-protein alignment (blastx) of the assembled *A. poculata* transcriptome was conducted against this database using BLAST v 2.6.0 (Altschul, Gish, Miller, Myers, & Lipman, 1990; Camacho et al., 2009). Transcripts with at least 80% identity and with at least 100bp mapped length were filtered from the transcriptome.

### Genome Annotation

To annotate repetitive content, *de novo* transposable element family identification and modeling was conducted using RepeatModeler v 1.0.1 (Smit & Hubley, 2008). RepeatMasker v 4.0.7 (A. Smit, Hubley, R & Green, P., 2015) was then used to soft-mask repetitive regions prior to gene modeling. Subsequent gene prediction included a multi-tool approach. First, *ab initio* gene prediction was done using GeneMark-ES v 3.51 (Lukashin & Borodovsky, 1998). Gene prediction using protein-based evidence was conducted with Exonerate v 2.4.0 (Slater & Birney, 2005) using the UniProt eukaryote database (downloaded December 28, 2017). RNA-seq reads from Chan et al. (2021) were aligned to the genome using STAR v 020201 (Dobin et al., 2013) and incorporated into the automated training of gene prediction using Braker2 v 2.1.2 (Brůna, Hoff, Lomsadze, Stanke, & Borodovsky, 2020). From the resulting predictions, high quality gene models (HiQ), here defined as those having >=90% RNAseq coverage support, were extracted. Gene prediction informed by transcriptomic evidence was carried out using PASA v 2.3.3 (Haas et al., 2003) with the flag --TRANSDECODER to keep only the longest open reading frame per group of transcript isoforms (see section ‘Transcriptome assembly’ above for transcriptome assembly details). Consensus gene predictions were acquired using EvidenceModeler v1.1.1 (Haas et al., 2008) weighting each line of evidence as follows: *ab initio* predictions: 1; RNA-seq based evidence: 1; protein-based evidence: 1; HiQ: 5; and transcriptomic-based evidence: 10. PASA v2.3.3 was run a second time to update gene models and add annotations of untranslated regions (UTRs) using the consensus gene prediction from EVidenceModeler. Finally, tRNAscan-Se v 1.3.1 (Chan & Lowe, 2019) was used to annotate transfer RNAs. Genome Annotation Generator (GAG) v 2.0.1 (Geib et al., 2018) was used to extract CDS, protein, and mRNA sequences.

Functional annotation of predicted genes was conducted based on sequence similarity using BLAST v 2.6.0 (Altschul et al., 1990; Camacho et al., 2009) searches via blastp with the settings of eval= 1e-5, max target seqs = 5, and max hsp = 1. Blast queries were conducted against three databases: NCBI nr, UniProt Swiss-Prot, and TrEMBL. Hits to each database were combined and annotated with Gene Ontology (GO) terms using the UniProt-GOA mapping.

### Investigating the possibility of whole-genome duplication

Given recent suggestion of a possible whole-genome duplication event in the genus *Acropora* (Mao & Satoh, 2019), we set out to determine whether a similar event may have occurred in *Astrangia poculata*. Detection and classification of duplication in the genome was carried out in several ways. First, estimates were conducted using the tool MCScanX under default settings (Wang et al., 2012). Results were then compared to a run under more relaxed settings (max gap size increased to 50). In both cases, duplication origins were classified using the duplicate_gene_classifier module.

A second method of whole-genome duplication (WGD) detection employed was the tool wgd v1.1.2 (Zwaenepoel & Van de Peer, 2018), which relies on the distributions of synonymous substitutions per synonymous site (*Ks*). To conduct the analysis, genome-wide coding sequences (CDS) were filtered for the longest translatable isoform of each CDS. wgd v1.1.2 was then run using the diamond aligner to compute the whole-paranome (the collection of all duplicate genes in a genome). A *Ks* distribution was constructed in pairwise mode and kernel density estimates were subsequently fit to the distribution and visualized. A *Ks* distribution for anchor pairs, defined as paralogs located on colinear duplicated segments, were similarly constructed and visualized. The shape of the *Ks* distribution was inspected for detection of ancient WGD with the expectation of an exponential decay shape in the absence of a WGD event (Zwaenepoel & Van de Peer, 2018).

Lastly, the whole-genome synteny aligner Satsuma2 (available at https://github.com/bioinfologics/satsuma2) was used to detect microhomologous regions between *A. poculata* chromosomes. Syntenic blocks of homologous sequences arranged in a colinear fashion between chromosomes were then then plotted using the tool Orthodotter (https://github.com/institut-de-genomique/orthodotter) to produce an Oxford grid (Edwards, 1991), an approach used previously to detect whole-genome duplication in arthropods (Schwager et al., 2017).

### Comparative genomics: Gene family and conserved synteny analysis

To characterize the genome organization and content of *Astrangia poculata* relative to other cnidarians, we completed several comparative genomic analyses. Phylogenetic orthology inferences were carried out using OrthoFinder2 v 2.4.0 (Emms & Kelly, 2018) on *Astrangia poculata* and 25 other available cnidarian proteomes using default parameters. Gene Ontology (GO) enrichment of gene families unique to *A. poculata* was conducted using the clusterProfiler package (Yu, Wang, Han, & He, 2012) implemented in R v 4.0.5 with a p-value cutoff of 0.05, a multiple testing correction method of Benjamini-Hochberg procedure (Benjamini & Hochberg, 1995), and a q-value cutoff of 0.2. GO enrichment analysis was conducted in the same manner for gene families that were shared amongst all cnidarians, representing a “core” cnidarian genome.

*Acropora millepora,* an obligately symbiotic coral of the complex clade, was selected for more detailed comparison with *Astrangia poculata* as its assembly was similarly complete. The sizes of gene families common to the two species were compared following the methods of González-Pech et al. (2021). Using the orthogroups previously identified by Orthofinder2, size differences of gene families shared between *A. poculata* and *A. millepora* were evaluated using Fisher’s exact test with the multiple testing correction method of Benjamini-Hochberg (Benjamini & Hochberg, 1995) and a significance threshold of adjusted *p* ≤ 0.05.

Conserved synteny between *Astrangia poculata* and *Acropora millepora* was assessed with MCScanX_h (Wang et al., 2012) using the homologous genes between the species identified by OrthoFinder2. Default parameters were used to identify colinear blocks (gap size of 25 genes allowable, minimum of 5 genes per colinear block).

## Results

### Genome statistics and assembly quality

Of the assemblers tried, wtdbg2 (Ruan & Li, 2020) produced the highest quality assembly (Table 1; Figure 2). After purging, polishing, and scaffolding, the assembly using wtdbg2 had an N_50_ of 31 Mb and a BUSCO score of 90.3% (89.1% single copies and 1.2% duplicate copies). The assembly size of 458 Mb is in line with BBMap predicted haploid size of 453 Mb (estimated ploidy of 2 and 40.95% repetitive). The 14 chromosome-level scaffolds match the known 2*n*=28 formula typical of most scleractinian corals (Figure 2A) (Flot et al., 2006). Further, the *k*-mer completeness was 52.24%, close to the expected 50% for a haploid assembly (Figure 2B). Gene prediction of the *A. poculata* genome assembly yielded 44,839 gene models. RepeatMasker predicted 39.29% of the genomic bases as repetitive elements, with an estimated GC content of 38.50%.

**Figure 2:**
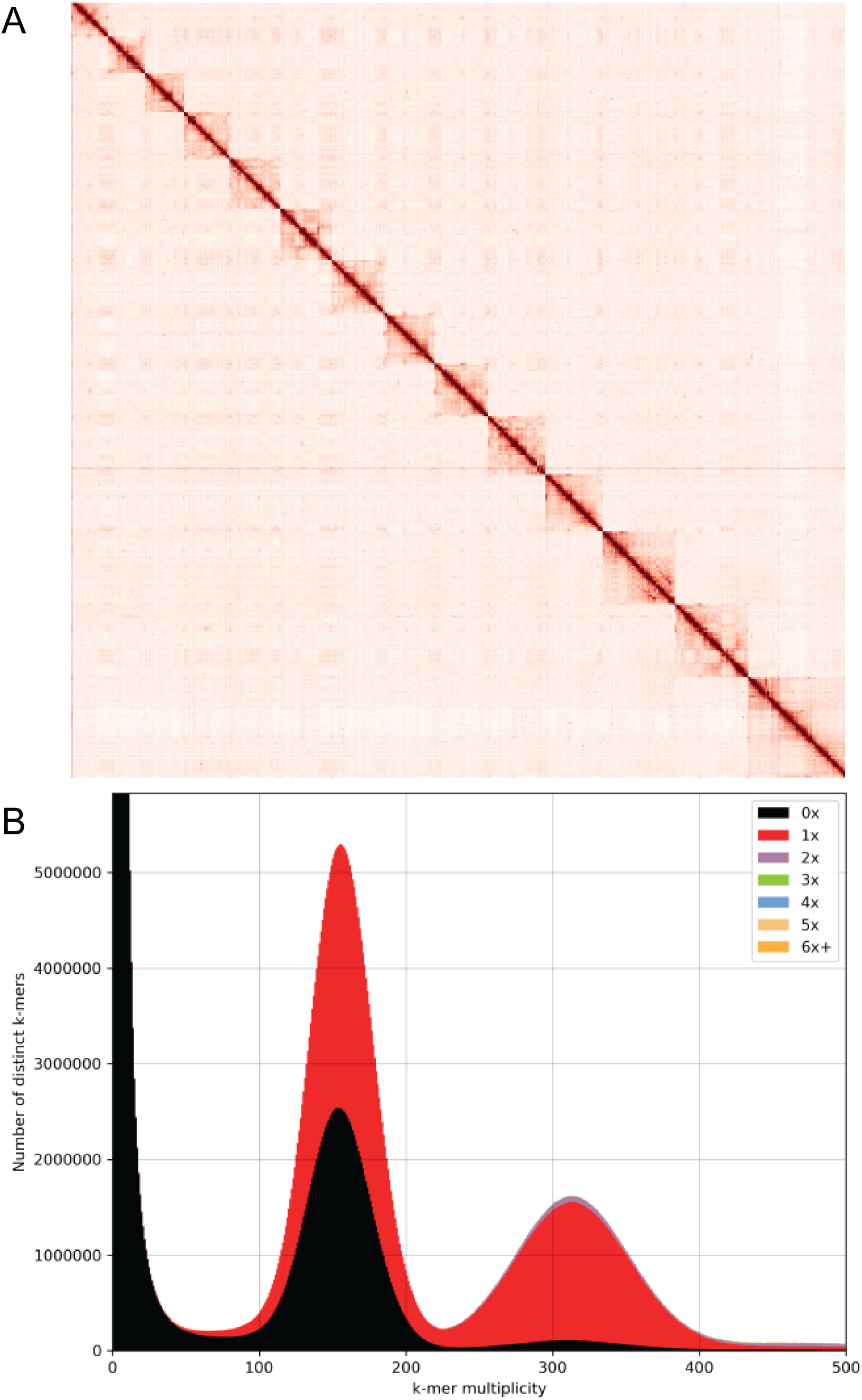
*Astrangia poculata* genome assembly quality. **A)** Proximity ligation sequencing data (Hi-C) contact map displaying the 14 chromosome-level scaffolds of the *A. poculata* assembly. Interaction points between chromosomes are represented by red dots with binning = 100 kb. Chromosomes are ordered by size from smallest to largest. **B)** KAT plot of distinct *k*-mer duplicity and the number of times these *k-* mers are represented in the final *A. poculata* genome assembly, with *k*=27.

**Table 1:** Summary of assembly statistics across assemblers. For each assembly strategy, information represents pre-purging, polishing, and scaffolding steps. Assembler references: ^1^ Vaser and Šikić (2019); ^2^Kolmogorov et al. (2019); ^3^Koren et al. (2017); ^4^Ruan and Li (2020).

| Assembler | Size (Mb) | Number of Contigs | N50 | BUSCO singles | BUSCO duplicated | K-mer completeness |
| --- | --- | --- | --- | --- | --- | --- |
| Ra<sup>1</sup> | 271 | 2,851 | 115 kb | 43.9% | 0.3% | 20.31% |
| Raven<sup>1</sup> | 400 | 2,912 | 172 kb | 52.3% | 0.4% | 27.19% |
| Flye<sup>2</sup> | 719 | 7,259 | 167 kb | 67.9% | 8.9% | 62.47% |
| Canu<sup>3</sup> | 597 | 8,317 | 97 kb | 56.0% | 7.9% | 41.75% |
| wtdbg2<sup>4</sup> | 476 | 4,423 | 439 kb | 55.1% | 0.3% | 36.47% |

### Whole-genome duplication

Duplication analysis via MCScanX (Wang et al., 2012) revealed 88 synteny blocks with 853 duplicated genes classified as putatively originating from whole genome or segmental duplication (Table 2). When relaxing MCScanX parameters by allowing a max gap size of 50 genes within colinear blocks (default gap size is 25), the number of duplications classified as whole genome or segmental increased to 1,005. Detected colinear blocks often involved more than two colinear regions, with some involving three or even four colinear regions. However, a *Ks* based approach using the tool wgd (Zwaenepoel & Van de Peer, 2018) resulted in an exponential decay shape of the distribution of synonymous substitutions per synonymous site, suggesting no signature of a WGD event in *A. poculata* (Figure 3). While we did detect many anchor pairs (colinear paralogs), the anchor *Ks* distribution also declines exponentially. This was consistent with the failure of Orthodotter to detect large colinear regions in the genome of *A. poculata* (Figure 4), suggesting that whole-genome duplication, if it occurred, was too ancient and the genome had subsequently undergone too much rearrangement to leave an obvious colinearity signature in the genome of *A. poculata*. Rather, these results suggest widespread duplications (tandem, proximal and dispersed) within the *A. poculata* genome, which may have resulted in novel functional gene copies through the process of neofunctionalization (Teshima & Innan, 2008).

**Figure 3:**
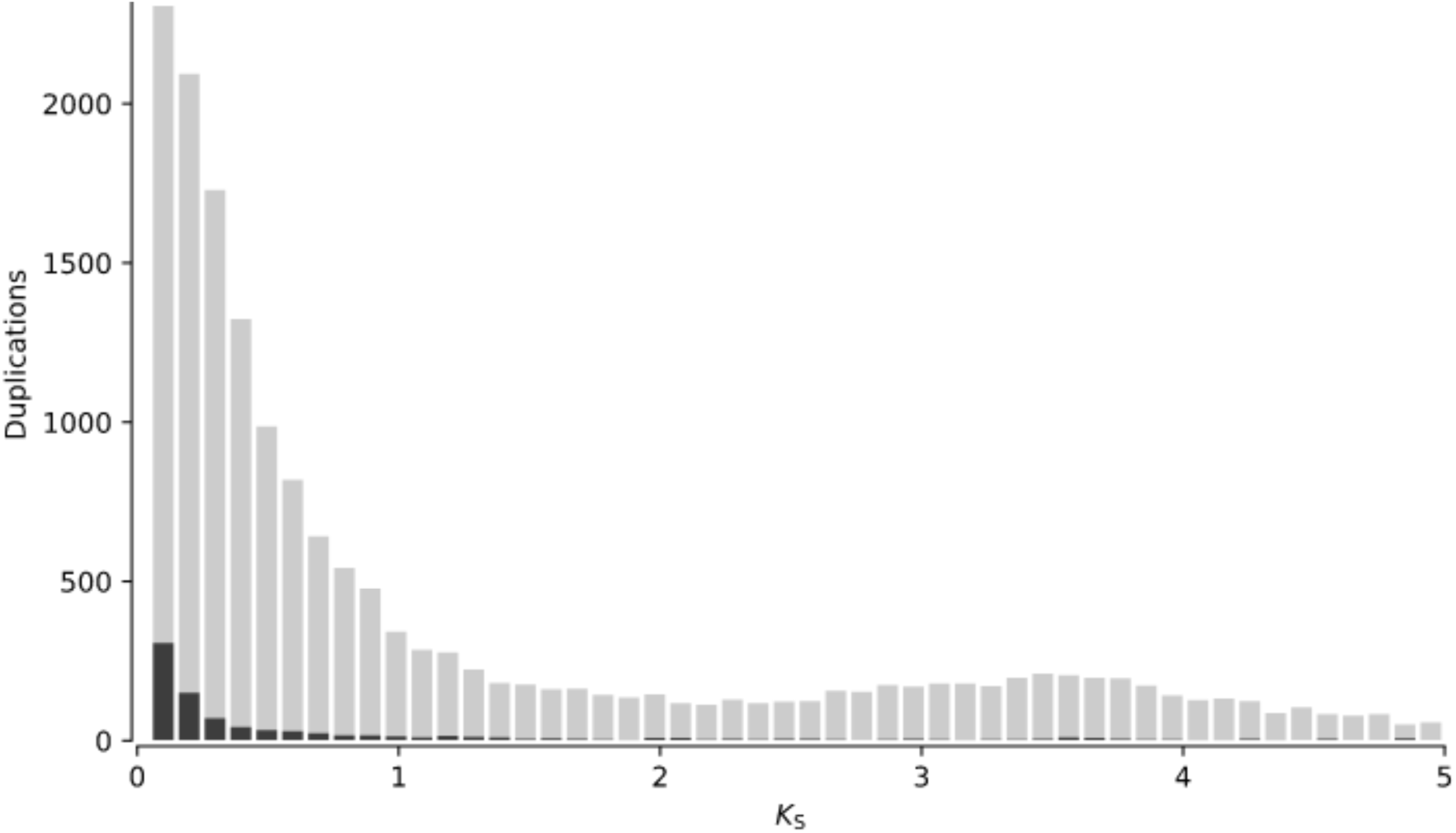
Distribution of synonymous substitutions per synonymous site (*Ks*) for all inferred duplications in the *A. poculata* genome with light grey representing all paralogous gene pairs and black representing anchor gene pairs.

**Figure 4:**
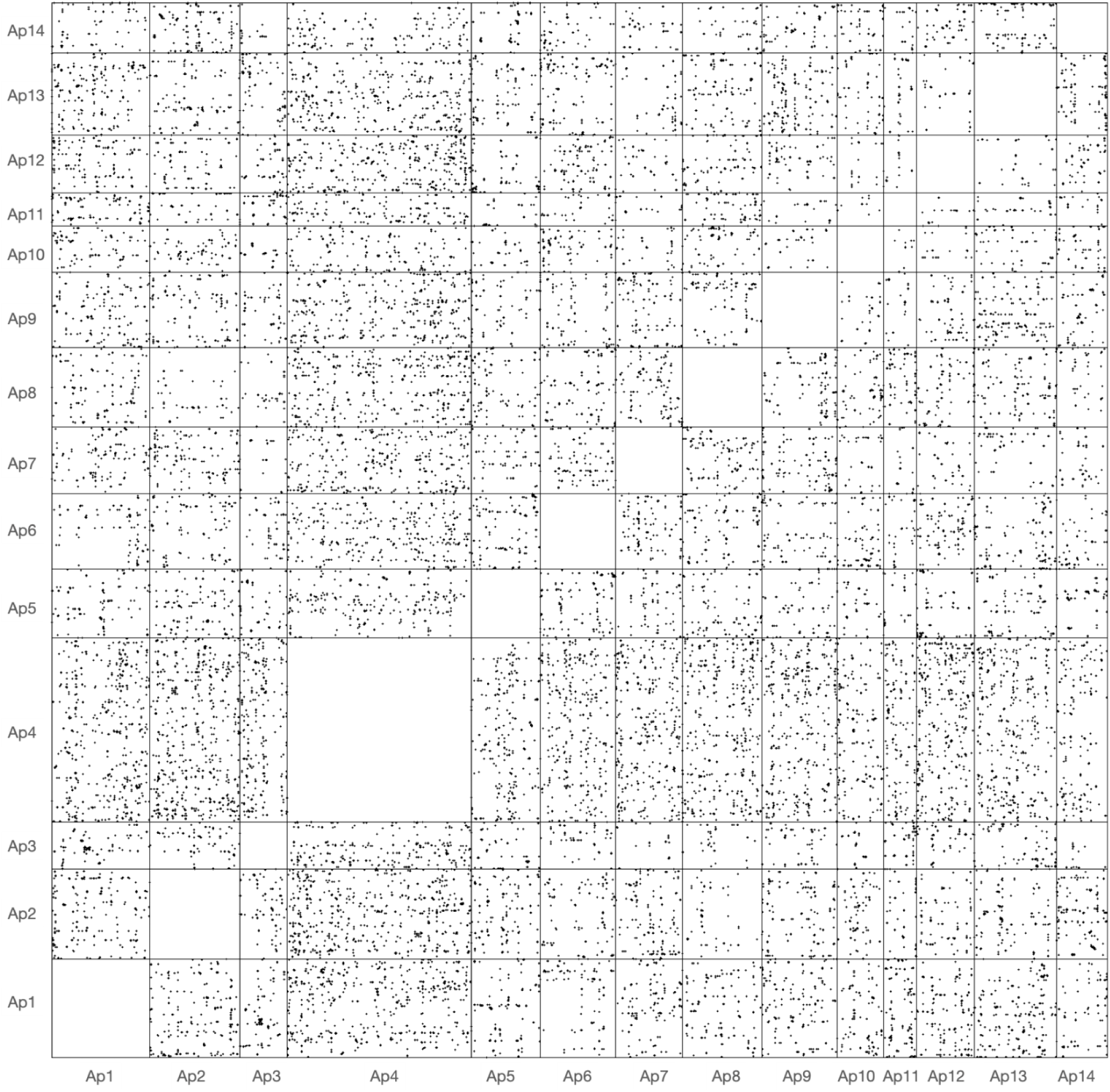
Oxford grid representing pairs of homologous regions detected by SatsumaSynteny across the genome of *A. poculata*. On this grid (not drawn to scale), each point represents a pair of identical or nearly identical 4096-bp regions. Ap1, Ap2… Ap14 represent the 14 chromosome-scale scaffolds in the assembly of the genome of *A. poculata*.

**Table 2:** Summary statics of duplication classification in the *A. poculata genome* with both default MCScanX parameters and with relaxed parameters that allow a gap of up to 50 genes.

| Type of duplication | Number with default parameters | Number with relaxed parameters |
| --- | --- | --- |
| Singleton | 11,063 | 11,063 |
| Dispersed | 20,794 | 20,702 |
| Proximal | 6,248 | 6,223 |
| Tandem | 8,198 | 8,163 |
| WGD or segmental | 853 | 1,005 |

### Comparative genomics: Gene family size and conserved synteny

Using a gene family analysis involving a total of 26 species, we defined a “core” cnidarian genome that consisted of 582 gene families shared among all cnidarians present in the analysis (Figure 5). A similar number of gene families (574) were found to be unique to *A. poculata.* Interestingly, only 179 orthogroups were present in all anthozoans included in the analysis. GO enrichment in clusterProfiler (Yu et al., 2012) of the gene families unique to *A. poculata* resulted in no significantly enriched GO terms (significance threshold of *p*<0.05 after adjusting for multiple testing). Likewise, gene families in the “core” cnidarian genome were not significantly enriched for any GO terms based on a similar GO enrichment test.

**Figure 5:**
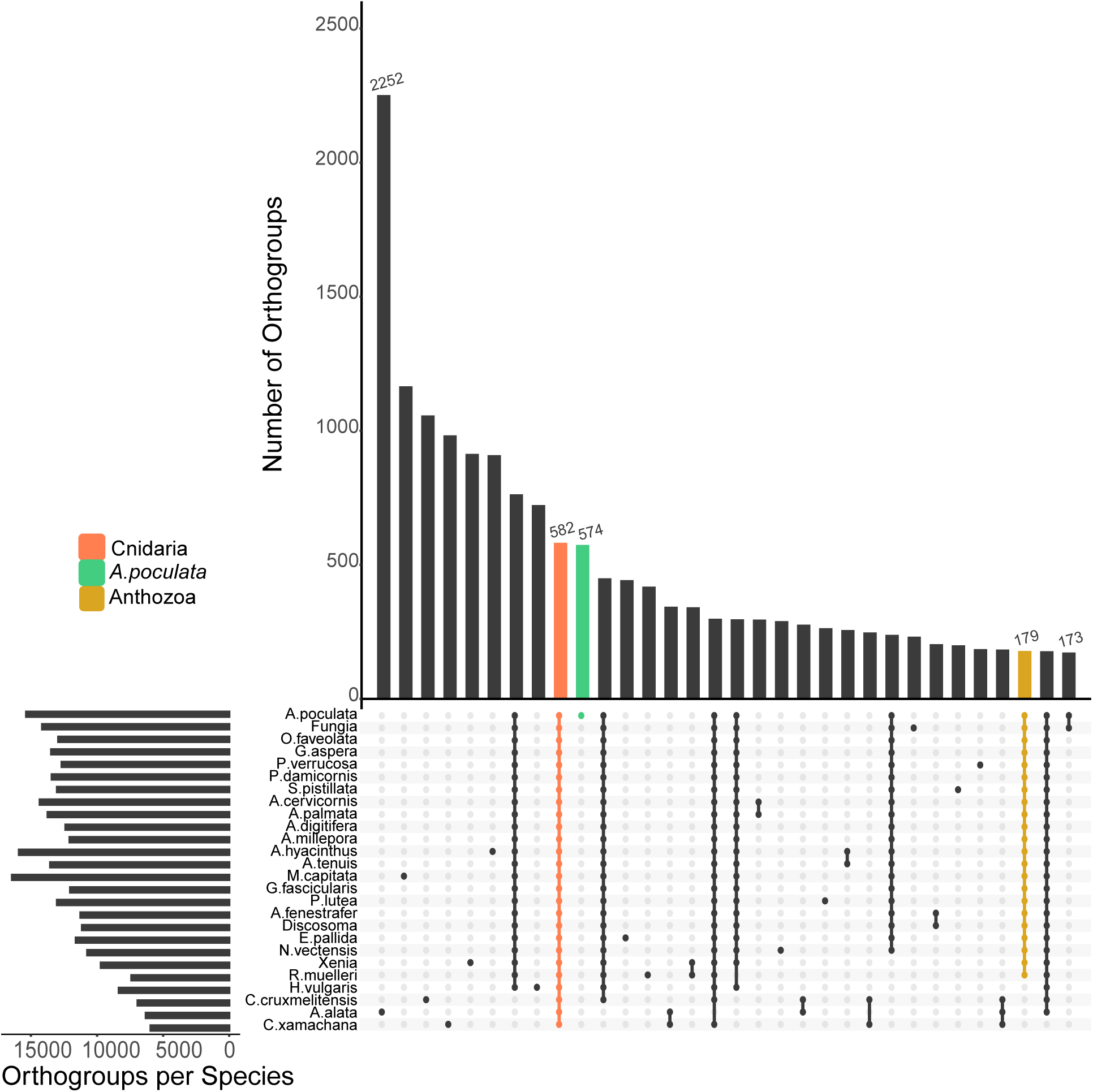
An UpSet plot representing the number of orthogroups containing each species included in the analysis. Each dot represents the presence of a given species in orthogroups, with orthogroups unique to *A. poculata* (green), shared amongst cnidarians (orange), and shared amongst anthozoans (gold).

Gene families that were common to *Astrangia poculata* and *Acropora millepora* were assessed for differences in gene numbers using Fisher’s exact test following the methods of González-Pech et al. (2021). In total, 165 gene families were identified as significantly different in gene numbers between the two species (Figure 6A; adjusted *p* ≤ 0.05). This included 71 gene families that were significantly larger in *A. millepora* and 94 gene families that were significantly larger in *A. poculata* (Figure 6A). Gene families larger in *A. millepora* that were most different in size compared to *A. poculata* (according to log_2_ fold change) included gene families that putatively encoded for zinc finger CCHC domain-containing proteins, cation channel sperm-associated proteins, RING-box proteins, lectins, and serine/threonine-protein kinases. In contrast, gene families larger in *A. poculata* that were most different in size compared to *A. millepora* by log_2_ fold change included orthogroups that putatively encoded for transposable elements, G protein-coupled receptors, Zinc finger MYM-type proteins, RNA-directed DNA polymerases, orexin receptors, ATP-dependent DNA helicases, E3 ubiquitin-protein ligases, and histone H3 (Figure 6B).

**Figure 6:**
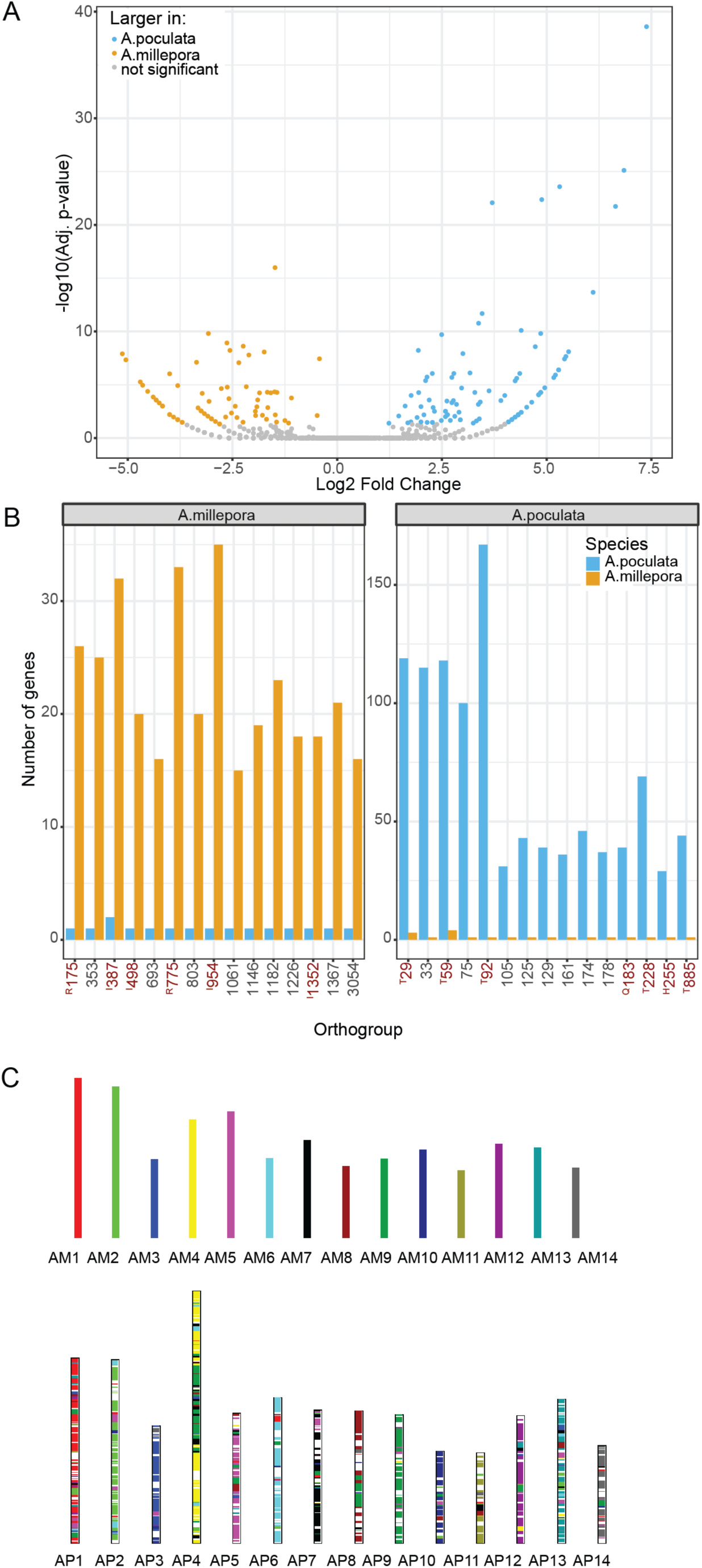
Comparison of *Astrangia poculata* and *Acropora millepora genomes.* **A)** Volcano plot of gene family size comparison using Fisher’s exact test between *A. poculata* and *A. millepora*. Points are colored according to whether they significantly larger in *A. poculata* (blue), significantly larger in *A. millepora* (gold), or not significant (grey) with adjusted *p* > 0.05. **B)** The top 15 (according to log_2_ fold change) significantly larger gene families in *A. poculata* (panel ‘A.poculata’) and *A. millepora* (panel ‘A.millepora’). Each bar represents the number of genes in the gene family for *A. poculata* (blue) and *A. millepora* (gold). Gene families with putative functions in reproduction, innate immunity, transposition, quiescence, and histone H3 are highlighted in red and designated with superscripts ‘R’ (reproduction), ‘I’ (immunity), ‘T’ (transposition), ‘Q’ (quiescence), and ‘H’ (histone H3). For readability, orthogroups are labeled with leading “OG” and zeros removed from their IDs. **C)** A bar plot representing the conserved synteny between the genomes of *Astrangia poculata* (AP) and *Acropora millepora* (AM). Each of *A. poculata’s* chromosomes are painted with the color of the *A. millepora* chromosome with which there is conserved synteny. White spaces indicate regions where colinear blocks were not detected.

In addition to gene family size analysis, orthologues common to *A. poculata* and *A. millepora* were evaluated for conserved synteny. MCScanX analysis of colinearity revealed a high level of conserved gene synteny between *A. poculata* and *A. millepora* with 3,451 syntenic blocks identified of at least 5 colinear genes. In total, 56.38% of orthologous genes were present in the colinear blocks between the two divergent coral species (Figure 6C).

## Discussion

*Astrangia poculata* has increasingly been used as a model coral system due to its temperature tolerance and flexibility in symbiont state (Aichelman et al., 2019; Burmester et al 2017; Burmester et al. 2018; Chan et al., 2021; Dimond & Carrington, 2007; Jacques et al., 1983; Peters, 1988; Sharp et al., 2017; DiRoberts et al. 2021; Wuitchik et al., 2021). Because *A. poculata* is facultatively symbiotic, the host and the algal symbiont response to manipulation can be distinguished—a study design that is often impossible in adult tropical corals, many of whom do not occur naturally in an aposymbiotic state. In addition to associating with an algal endosymbiont of the family Symbiodiniaceae, *A. poculata* creates a calcium carbonate skeleton similar to reef-building corals (Hayes & Goreau, 1977; Peters, 1988). However, corals are a diverse group of organisms with respect to their biology and ecology (e.g., habitat, morphology, and response to environmental change) (Chappell, 1980; Fabricius et al., 2011; Kusumoto et al., 2020; Muir, Wallace, Pichon, & Bongaerts, 2018). Thus, it is important to recognize the differences, as well as the similarities, between *A. poculata* and other corals. Here, we have developed a chromosome-scale genome assembly for *A. poculata*, which has allowed us to characterize some of these differences and similarities between *A. poculata* and other cnidarians. These results have revealed insight into potential genomic drivers of cnidarian biology, such as *A. poculata’s* astounding ability to enter a dormancy state when exposed to extreme cold temperatures, and elucidated the demands placed on sperm during mass spawning events of tropical corals.

### No evidence of ancient whole genome duplication, but recent duplications are abundant

Whole-genome duplication (WGD) can provide new genetic material upon which selection or genetic drift may act (Holland & Ocampo Daza, 2018; Ohno, 1970). A possible WGD event in the most recent common ancestor of the genus *Acropora* was suggested using phylogenomic and comparative genomic techniques (Mao & Satoh, 2019). However, it is unknown if such an event may have occurred at other points in the scleractinian lineage where polyploidism is common. Our conservative results from MCScanX indicated that 853 duplicated genes (1.8%) were classified as possibly originating from large-scale duplication events, such as segmental or WGD. To further identify whether this may indeed be indicative of an ancient WGD event, we examined the distribution of synonymous substitutions per synonymous site (*Ks*). Under a model of a constant rate of duplication and loss, there should be an exponential decay shape to a *Ks* distribution, which is the shape we find in the distribution for *A. poculata* (Figure 3). In contrast, when a WGD event has occurred, it leaves a signature peak in the *Ks* distribution (Zwaenepoel & Van de Peer, 2018). Additionally, we examined the *Ks* distribution of the anchor pairs (paralogs located on colinear duplicated segments). However, many anchors detected represented small *Ks* values (0-0.1) and also followed an exponentially decaying shape. Similarly, a search for microsynteny using SatsumaSynteny followed with colinearity detection using Orthodotter did not reveal abundant pairs of colinear regions (Figure 4). These results suggest that while large scale segmental duplications may be present in the *A. poculata* genome, we are unable to detect a strong signature of an ancient WGD event. Because these approaches may not be able to detect very ancient duplications, a phylogenomic approach (Zwaenepoel & Van de Peer, 2018) will be required to further test for an ancient whole-genome duplication event in Scleractinia as high-quality genome assemblies representative of each taxonomic group across the cnidarian phylogeny become available.

### Genome content of temperate *Astrangia poculata* versus tropical *Acropora millepora*

Though the orthologue analysis resulted in 574 orthogroups unique to *A. poculata*, no significantly overrepresented GO terms were present in these gene families (Figure 5). This could be partially explained by the sensitivity of such analyses to large differences in genome assembly quality (Liu, Hunt, & Tsai, 2018). The *A. poculata* genome assembly is among the most complete and contiguous coral genomes to date. Many of the other currently available cnidarian genome assemblies remain considerably more fragmented. For this reason, we limited our subsequent analyses to comparisons between *Astrangia poculata* and *Acropora millepora*. *A. millepora* has a chromosome-scale genome assembly (Fuller et al., 2020), and contrasts with *A. poculata* in its ecology and evolutionary history. While *A. poculata* is a facultatively symbiotic temperate coral of the robust clade, *A. millepora* is an obligately symbiotic tropical coral of the complex clade.

### Conserved microsynteny between complex and robust clades

Synteny analysis between *A. poculata* and *A. millepora* revealed considerable conserved colinearity (56.38%; Figure 6C) despite approximately 415 Mya of divergence of the two clades, Robusta (*A. poculata*) and Complexa (*A. millepora*) (Stolarski et al., 2011). This surprisingly high level of colinearity is in line with previous work comparing other complex and robust coral species. Ying et al. (2018) found that the extent of conserved gene order within Scleractinia, regardless of clade, was relatively high compared to the level of conserved synteny between sea anemones *Exaiptasia* and *Nematostella* in the order Actinaria. Our results further lend support to this conclusion of consistently high conserved gene order across scleractinians.

### Differential gene family expansions related to innate immunity and symbiosis

While gene order analysis highlighted similarities between *A. poculata* and *A. millepora*, gene family size comparisons revealed differences (Figure 6A; Figure 6B). Gene family expansions are often observed during adaptation in corals (van Oppen & Medina, 2020). Notably, of the gene families expanded in *A. millepora* relative to *A. poculata*, the gene family with the largest log_2_ fold change contained several copies of Zinc finger CCHC domain-containing protein 3, which plays a role in innate immune response to viruses (Lian et al., 2018). Defense against viruses is likely important in *A. millepora,* as previous work has identified massive viral outbreaks in this species (Correa et al., 2016). Further, obligately symbiotic cnidarians have a more advanced innate immunity repertoire relative to non-symbiotic relatives, possibly driven by the dynamic and constant interaction with the algal endosymbiont (Cunning, Bay, Gillette, Baker, & Traylor-Knowles, 2018; Shinzato et al., 2011; Shumaker et al., 2019; van Oppen & Medina, 2020; Voolstra et al., 2017). These previous findings relied on comparisons to more distantly-related, non-symbiotic anthozoans. However, here we determine that this holds with a comparison to a more closely related facultatively symbiotic coral, as opposed to a non-coral anthozoan relative.

The molecular elements governing the uptake, maintenance, and breakdown of symbiosis in corals still remain largely unclear. However, previous work indicates that features of the innate immune system of the host, notably lectins, play an important role (Fransolet, Roberty, & Plumier, 2012; Hu, Zheng, Fan, & Zheng, 2020; Kvennefors, Leggat, Hoegh-Guldberg, Degnan, & Barnes, 2008; Takeuchi et al., 2021; Zhou et al., 2018). Lectins are pattern recognition proteins that bind to carbohydrates (Goldstein, Hughes, Monsigny, Osawa, & Sharon, 1980). Within the top 15 orthogroups significantly larger in *A. millepora* relative to *A. poculata* were gene families related to innate immunity that have previously been implicated in establishing symbiotic associations in corals, including C-type lectins and macrophage mannose receptors that mediate endocytosis of glycoproteins (Figure 6B; orthogroups OG0000498, OG0001352, and OG0000387). Kvennefors et al. (2008) isolated a mannose-binding lectin in *A. millepora* and demonstrated its affinity to binding to both pathogens and algal dinoflagellates of the family Symbiodiniaceae. Since then, comparative genomics and single-cell RNA sequencing have further emphasized the role of lectins in the cnidarian-algal symbiosis in additional species (Cunning et al., 2018; Hu et al., 2020). Here, we were able to compare two scleractinian corals, one facultatively symbiotic and the other obligately symbiotic. We found that pattern recognition proteins (e.g., lectins) are expanded in *A. millepora* relative to *A. poculata*. Future study characterizing the potential differences in the establishment and maintenance of symbiosis between obligately vs facultatively symbiotic corals is warranted, and the results here highlight lectin-related gene families as excellent targets.

### Sexual reproduction in mass spawning tropical corals

In addition to innate immunity, top gene families larger in *A. millepora* compared to *A. poculata* included those related to sexual reproduction (Figure 6B), with functions involved in sperm cell hyperactivation (cation channel sperm-associated protein subunit epsilon; orthogroup OG0000775) and meiosis (RING-box protein 1; orthogroup OG0000175). These findings are interesting because of the difference in reproductive modes between *A. millepora* and *A. poculata*. *Astrangia poculata* is a gonochoric species that reproduces via broadcast spawning wherein gametes are released into the water column prior to fertilization (Szmant, Yevich, & Pilson, 1980). Spawning occurs annually from August to September based on the seasonal maximum temperature, with a second cycle sometimes observed in October or November (Peters, 1988). *A. poculata* sperm are unlikely to have to compete with many other corals for fertilization of eggs because there are only a few other temperate coral species. In contrast, *A. millepora* is a hermaphroditic broadcast spawning species that reproduces during ‘mass spawning’ events synchronized with the occurrence of the full moon (Kaniewska, Alon, Karako-Lampert, Hoegh-Guldberg, & Levy, 2015). During these annual spawning events, *A. millepora* releases gametes simultaneously with over 100 other coral species, as well as hundreds of other invertebrates over the course of only a few nights (Babcock et al., 1986; Harrison, 2011). This creates competition between gametes, as well as the opportunity for interspecific hybridization. Expansion of these reproductive-related gene families have the potential to soften or maintain species boundaries and would be great targets for future studies examining adaptation to mass spawning in corals.

### Genome plasticity in *A. poculata*: transposition and epigenetics

Gene families that were larger in *Astrangia poculata* relative to *Acropora millepora* included several families of transposable elements (Figure 6B; orthogroups OG0000092, OG0000228, OG0000885, OG0000029, and OG0000059). Retrotransposition has been found to contribute to gene family expansion in Symbiodiniaceae (González-Pech et al., 2021; Lin et al., 2015) and, based on our results, may also play a role in the host *A. poculata*. Further, transposable elements promote adaptation and drive genome plasticity in many species, including bacteria, fungi, plants, and animals (Bennett, 2004; Feschotte & Pritham, 2007; Leitch & Leitch, 2008; Mat Razali, Cheah, & Nadarajah, 2019; Yuan et al., 2021). Epigenetic modification is involved in host genome regulation of transposable elements (Matzke, Mette, & Matzke, 2000). Interestingly, we also found that a family of histone H3 proteins was expanded in *A. poculata* relative to *A. millepora* (Figure 6B; orthogroup OG0000255). In eukaryotes, histone H3 is one of the core histone proteins involved in structuring chromatin (Kornberg, 1977; McGhee & Felsenfeld, 1980). The sequence variants, as well as different modification states of histone H3 are thought to influence gene regulation (Hake et al., 2006; Jiang & Berger, 2017; Klemm, Shipony, & Greenleaf, 2019; Kouzarides, 2007; Loyola & Almouzni, 2007; Maehara et al., 2015; Sarma & Reinberg, 2005). In plants, histone H3 plays a role in development and abiotic stress (Otero, Desvoyes, & Gutierrez, 2014; Yuan, Liu, Luo, Yang, & Wu, 2013). These results suggest that transposable elements and epigenetic modification may play an important role in the plasticity of *A. poculata*.

### Winter quiescence in A. poculata

Among the top gene families expanded in *A. poculata* was a family of G-coupled protein receptors (orthogroup OG0000183), including RYamide receptors, orexin receptors, and neuropeptide SIFamide receptors (Figure 6B). In *Drosophila*, RYamide receptors are possibly associated with feeding suppression (Ida et al., 2011), while neuropeptide SIFamide receptors have been associated with promotion of sleep (Park, Sonn, Oh, Lim, & Choe, 2014). Similarly, orexin receptors are known to regulate circadian sleep/wake cycles in mammals (Chemelli et al., 1999). The expansion of this gene family in *A. poculata* relative to *A. millepora* may explain *A. poculata*’s ability to enter a dormant state when exposed to near freezing temperatures. During winter months in the intertidal and subtidal regions at the northernmost edges of the species’ range, *A. poculata* has been demonstrated to enter this dormancy, referred to as “winter quiescence”, when water temperatures plummet to below 6°C (Grace, 2017). During this state, polyps are retracted, and oral plates are puffed out, while feeding is reduced or ceased entirely (Grace, 2017; Jacques et al., 1983) (Figure 1B). This finding has relevance to tropical systems, in which some Mediterranean anthozoans have been shown to enter a similar “summer dormancy” state, including corals (Caroselli, Falini, Goffredo, Dubinsky, & Levy, 2015; Coma, Ribes, Gili, & Zabala, 2000). Overall, these results indicate that gene family expansions may have contributed to adaption of *A. poculata* to the high variance in environmental conditions that this species experiences temporally and spatially across its range.

## Conclusion

In this study, we present the first chromosome-scale assembly of the facultatively symbiotic, temperate coral *Astrangia poculata*. Our contribution of a high-quality genome resource for *A. poculata* advances the use of this species as an experimental model and lays the groundwork for numerous future studies, as the *A. poculata* is an important emerging model for coral health (Neff, 2020). Further, comparison of the *A. poculata* genome to the tropical, obligately symbiotic coral *Acropora millepora* uncovered potential genomic drivers of unique features of not only *A. poculata*, but of *A. millepora*, as well. Taken together, these results have generated genomic targets for future study of adaptation in these species and emphasize the power of comparative genomics to reveal novel insights into the biology of corals.

## Acknowledgements

We thank all authors who made their coral genome assemblies available for the comparative genomic analyses in this work. This work was made possible by NSF grant OCE-1537959 to IBB, NIH grant T32: Computation, Bioinformatics, and Statistics (CBIOS) Training Program to KHS, and the Pennsylvania State University Biology Department. Special thanks to Sam Piorkowski and Claire Klippel for their computational and aquaria support.

